# Resting-state hyper- and hypo-connectivity in early schizophrenia: which tip of the iceberg should we focus on?

**DOI:** 10.1101/2024.09.20.613853

**Authors:** David Tomecek, Marian Kolenic, Barbora Rehak Buckova, Jaroslav Tintera, Filip Spaniel, Jiri Horacek, Jaroslav Hlinka

## Abstract

In this study, we explore the intricate landscape of brain connectivity in the early stages of schizophrenia, focusing on the patterns of hyper- and hypoconnectivity. Despite existing literature’s support for altered functional connectivity (FC) in schizophrenia, inconsistencies and controversies persist regarding specific dysconnections.

Leveraging a large sample of 100 first-episode schizophrenia patients (42 females/58 males) and 90 healthy controls (50 females/40 males), we compare the functional connectivity across 90 brain regions of the Automated Anatomical Labeling atlas. We inspected the effects of medication and examined the association between FC changes and duration of untreated psychosis, duration of antipsychotic treatment, as well as symptom severity of the disorder. Our approach also includes a comparative analysis of three denoising strategies for functional magnetic resonance imaging data.

In patients, 15 region pairs exhibited increased FC, whereas 150 pairs showed reduced FC relative to controls. Despite this numerical asymmetry, the overall distribution of FC changes was relatively balanced: the median FC was not systematically shifted, indicating no global tendency toward either hyper- or hypoconnectivity. Notably, seveFC alterations were significantly associated with variability in symptom severity and antipsychotic medication across patients.

Taken together, these results suggest a pattern of localized dysconnections embedded within an otherwise globally balanced change in connectivity profile in early schizophrenia. Importantly, this balance was substantially disrupted towards dominant observation of hypoconnectivity when less stringent denoising strategies were applied, with results increasingly dominated by hypoconnectivity, pointing to data preprocessing as a critical source of variability across studies.

## 1 Introduction

Although a neurodevelopmental hypothesis for schizophrenia complemented by neurodegenerative processes following the onset of psychosis is now well established, it is not clear how these processes are linked to the brain dysfunction underlying specific schizophrenia psychopathology (Dauvermann et al. 2017). The converging evidence from brain imaging studies indicates that these processes lead to disordered brain inter-regional functional connectivity (FC), and dysconnection represents the major candidate for the pathophysiological substrate of psychotic symptoms (Friston 2002). Concretely, the dysconnection concept suggests that the specific symptoms of schizophrenia can be described in terms of reduced or increased functional coupling between distinct brain regions (Friston & Frith 1995; Stephan et al. 2006). This concept is in line with earlier intuition referring to the title of the disease (i.e., “fragmented mind”), and it has been conceptualized in a neurobiological framework proposing a disruption of the anatomical and functional connectivity between brain areas as the neurobiological correlate of altered information processing (Friston & Frith 1995). To quantify such connectivity disruption, the concept of functional connectivity, formalized as temporal covariation of neural signals between spatially disparate brain areas (Friston et al. 1993), is often used.

The functional dysconnection in schizophrenia has been reported to affect connections of a multitude of brain regions, such as the frontal lobe, including language areas (Backash et al. 2014; Benetti et al. 2009; Cole et al. 2011; Weinberger et al. 1994), sensory-motor cortex (Kaufmann et al. 2015), temporal and limbic structures (Crossley et al. 2009; Meyer-Lindenberg et al. 2005), and thalamus (Anticevic et al. 2014; Giraldo-Chica & Woodward 2017; Guller et al. 2012; Woodward & Heckers 2016; Woodward et al. 2012). Despite the fact that neuroimaging studies strongly support the role of altered FC in schizophrenia, these reported findings are highly inconsistent; specific regions associated with disconnection in schizophrenia still remain controversial, and a robust conclusion has not yet been obtained. Some of the probable reasons for these inconclusive results are the heterogeneity of clinical samples regarding the stage of illness (T. Li et al. 2016) and low sample sizes based on up to 40 participants on average (Dong et al. 2018). Indeed, a small sample size has been previously systematically documented to give rise to heterogeneous localized findings of white matter abnormalities in first-episode psychosis (Melicher et al. 2015). We further conjecture that variability in the data preprocessing and analysis approaches might also contribute to the variability of the observed results (Burgess et al. 2016). The majority of previous rs-fMRI studies in schizophrenia reported hypoconnectivity between brain regions (Dong et al. 2018; S. Li et al. 2019), but hyperconnectivity has also been repeatedly observed in various disease stages (T. Li et al. 2016; Dong et al. 2018; González-Vivas et al. 2019; Anticevic et al. 2015). Longitudinal studies of FES subjects reported (partial) normalization in terms of increased fMRI activation after antipsychotic treatment (González-Vivas et al. 2019). Another factor responsible for the heterogeneity of the findings is the methodological approaches focused on FC between a priori preselected regions, as contrasted with a whole-brain analysis based on the parcellation according to suitable anatomical or functional brain atlases. Given these principal differences, the reported findings are strongly influenced by the a priori choice of the regions of interest, which in some cases may not include all the pathophysiologically relevant areas. On the other hand, whole-brain studies may suffer from the fact that the correction for multiple comparisons could bring false negative results, specifically in the case of underpowered studies. Our study aimed to remediate some of the above-mentioned inconclusiveness and to utilize the whole-brain FC analysis on a large cohort of 100 first-episode schizophrenia (FES) patients, ensuring high homogeneity in terms of chronicity and exposure to medication. The connectivity was calculated between all 90 brain regions comprising the Automated Anatomical Labeling (AAL) template image (Tzourio-Mazoyer et al. 2002) and compared with a group of 90 healthy controls. On the basis of the previous evidence, we hypothesized that schizophrenia would show decreased connectivity between the candidate brain regions. Furthermore, we tested whether FC changes were moderated by the effect of medication and whether they were linked to clinical factors such as positive and negative symptoms of schizophrenia. In addition, we evaluated two different denoising strategies of the fMRI data, as this step potentially affects the resulting FC measures (Burgess et al. 2016).

## 2 Methods

### 2.1 Study overview and samples

In total, 190 subjects participated in the study: 100 FES patients (mean age=28.78, SD=6.84, 42 females/58 males) and 90 healthy volunteers serving as controls (mean age=27.88, SD=6.83, 50 females/40 males). There were no significant differences between the patient and control samples in age and sex. In the patient group, at the time of the MRI scan, the average duration of untreated psychosis was 3.23 months (SD = 4.79), and the average duration of antipsychotic treatment was 2.29 months (SD = 4.55). The study design was approved by the local Ethics Committee of the Institute of Clinical and Experimental Medicine and the Psychiatric Center Prague. All subjects provided written informed consent after receiving a complete description of the study.

The FES patients were diagnosed according to ICD-10 criteria and structured MINI International Neuropsychiatric Interview (Sheehan et al. 1998). FES subjects were investigated during their first hospitalization in the Prague psychiatric hospitals with a catchment area of 1 million inhabitants. Patients were considered as FES if they fulfilled these criteria: a) first hospitalization for schizophrenia, and b) clinical interview identified first psychotic and/or prodromal symptoms of psychosis not earlier than 24 months ago (mean=5.90 months, SD=6.16).

The resting fMRI was performed at the initial stage of second-generation antipsychotic therapy (mean 10 weeks of medication at the time of rsfMRI). The mean dose of chlorpromazine equivalents (Woods 2003) was 388.87 mg (SD=235.44) per day. Psychometrics included the Positive and Negative Symptom Scale (PANSS) (Kay et al. 1987). Ninety healthy control subjects (HC) were recruited via a local advertisement; they had a similar socio-demographic background as the FES to whom they were matched by age and sex.

The HC had a slightly higher number of years of education than the FES (HC: mean=16.56, SD=2.33; FES: mean=13.51, SD=2.46; t=7.033, p<0.001). HC were evaluated with MINI (Sheehan et al. 1998) and were excluded if they had a lifetime history of any psychiatric disorder or a family history of psychotic disorders. Other exclusion criteria for both groups included a history of seizures or significant head trauma, mental retardation, a history of substance dependence, and any MRI contraindications. Written informed consent was obtained from all participants.

### 2.2 fMRI data acquisition

Scanning was performed using a 3T Siemens Magnetom Trio MRI scanner located at the Institute of Clinical and Experimental Medicine in Prague, Czech Republic. Functional images were obtained using T2*-weighted gradient echo-planar imaging (GR-EPI) sequence with blood oxygenation level-dependent (BOLD) contrast with the following parameters: repetition time (TR)=2000 ms, echo time (TE)=30 ms, flip angle=70°, voxel size=3×3×3 mm, matrix size 48×64 voxels, 35 axial slices acquired continuously in sequential decreasing order covering the entire cerebrum, 400 functional volumes in total. Also, for anatomical reference, a three-dimensional T1-weighted high-resolution magnetization-prepared rapid gradient echo (MPRAGE) image was acquired from each participant using the following parameters: repetition time (TR)=2300 ms, echo time (TE)=4.63 ms, flip angle=10°, voxel size=1×1×1 mm, matrix size 256×256 voxels, 224 sagittal slices covering the entire brain.

### 2.3 Data preprocessing, brain parcellation, and FC analysis

Functional MRI is a neuroimaging method that is based on measuring blood oxygen level-dependent (BOLD) signal (Ulmer & Jansen 2020). One of the typical features of the fMRI data is the noise that is present in the raw BOLD signal (Biswal et al. 1995; Raichle 2015), which significantly limits the reliability of functional connectivity measures (Shirer et al. 2015). Typical artifacts, such as subject movements, arterial pulsation, respiration, and also the hardware of the MRI scanner itself, induce non-neural temporal correlations in the BOLD (Pamilo et al. 2015), and relatively sophisticated data preprocessing is warranted to maximize the level to which the functional connectivity estimates reflect the underlying neuronal dynamics.

First, the structural and functional images were converted from DICOM to NIFTI format using the dcm2niix tool (X. Li et al. 2016). The following preprocessing steps were performed with the functional data: functional realignment and unwarp; slice-timing correction; outlier identification; direct segmentation and normalization; and functional smoothing - pipeline labeled as "default preprocessing pipeline for volume-based analyses (direct normalization to MNI-space)" in CONN toolbox (Whitfield-Gabrieli & Nieto-Castanon 2012).

The denoising steps included the regression of six head-motion parameters with their first-order temporal derivatives and five principal components of WM and CSF signals using a component-based noise correction method (CompCor) implemented in the CONN toolbox (Behzadi et al. 2007). Time series from defined regions of interest were additionally linearly detrended and finally filtered by a band-pass filter with cutoff frequencies 0.008-0.09 Hz. We shall refer to this preprocessing setup as the *stringent* denoising scheme.

As an alternative denoising pipeline, closer to the practice in some studies (Argyelan et al. 2014; Wang et al. 2015), we used a *moderate* denoising scheme including six head-motion parameters without their first-order derivatives and only the mean time series of WM and CSF. This pipeline was performed without explicit linear detrending. However, time series were also finally filtered by a band-pass filter with cutoff frequencies 0.008-0.09 Hz.

To isolate the impact of denoising, we conducted another analysis on the data that underwent the same default preprocessing pipeline in the CONN toolbox (Whitfield-Gabrieli & Nieto-Castanon 2012) as described above, while explicitly omitting all denoising procedures. This variant is referred to as the *raw*.

### 2.4 Analysis

For each subject, to quantify the whole-brain pattern of functional connectivity, we performed an ROI-to-ROI connectivity analysis by computing Pearson’s correlation matrix among the regional mean time series extracted from 90 brain regions (excluding the cerebellar regions) of the Automated Anatomical Labeling (AAL) atlas (Tzourio-Mazoyer et al. 2002) using the CONN toolbox (Whitfield-Gabrieli & Nieto-Castanon 2012). The resulting connectivity matrices were represented as Fisher’s z-transformed Pearson’s correlation coefficients. Although participants were balanced in terms of age and sex, these variables were further included as nuisance covariates in the linear regression model to further minimize their potential influence on functional connectivity.

We used a two-sample Mann-Whitney U-test to evaluate differences between groups. Spearman’s correlation was used to determine the relation between the four subscales of the PANSS scale (Kay et al. 1987), antipsychotic medication at the time of the MRI scan (measured by chlorpromazine equivalent), duration of untreated psychosis, duration of antipsychotic treatment, and functional connectivity in patients. To increase the statistical analysis power, these analyses were limited to ROI pairs that showed a significant effect of disease (i.e., effect in the initial between-group comparison). The resulting p-values from each analysis were corrected for multiple testing by controlling the false discovery rate (FDR) (Benjamini & Hochberg 1995). Additional analyses with the moderate and raw denoising schemes were performed using the same analytical pipeline.

## 3 Results

### 3.1 The difference between healthy controls and schizophrenia patients

In the group comparison using the default, stringent denoising scheme, a difference was observed in a multitude of ROI pairs. 150 ROI pairs had significantly greater functional connectivity (FDR-corrected p*<*0.05) in patients, while healthy volunteers exhibited significantly greater (FDR-corrected p*<*0.05) functional connectivity in 15 ROI pairs compared to patients (Figure 1). Despite the numerical prevalence of observed hyperconnections in patients, healthy volunteers, and patients did not show a difference in the median functional connectivity (median FC in patients=0.0709, MAD=0.0182, median FC in controls=0.0653, MAD=0.0209, u=4007 (z-score=-0.8663), p=0.1932).

**Figure 1:**
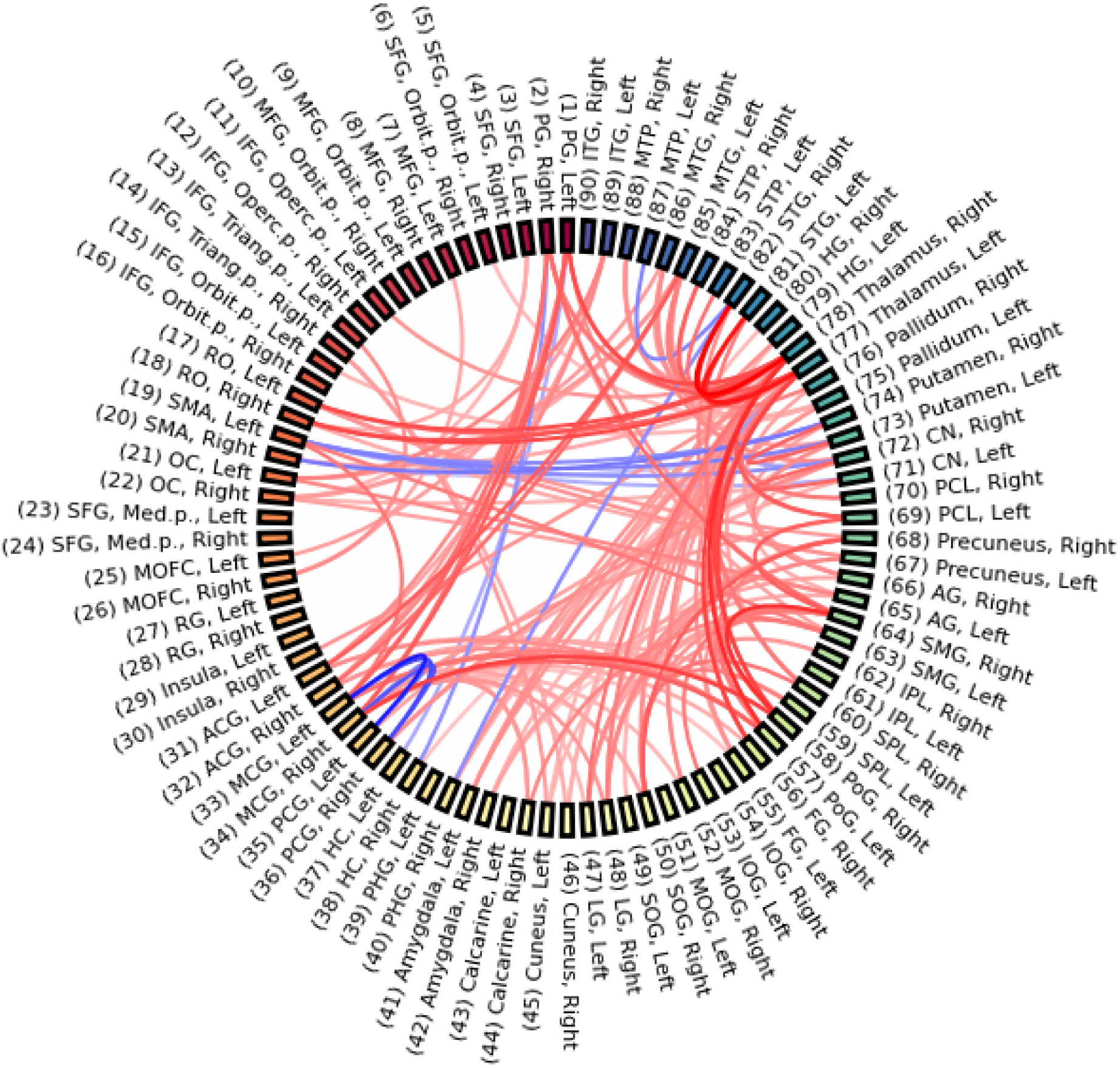
Difference between HC and FES; higher FC in healthy controls = blue, higher FC in patients = red, p*<*0.05 (FDR-corrected).

The most increased links in patients included a range of connections involving particularly the bilateral thalamus, right middle cingulate gyrus, right supramarginal gyrus, left middle temporal gyrus (see Table S1 for the full list of affected connections). In the opposite direction, the decreased links in patients comprised especially the left supplementary motor area, bilateral posterior cingulate gyrus, right superior temporal gyrus, and left middle cingulate gyrus (see Table S2 for the full list of affected connections). See Figure 2 for a graphical overview of the group comparison of healthy controls and patients, including its stability with respect to additional control for inter-subject variability in the amount of head motion; see Discussion for details.

**Figure 2:**
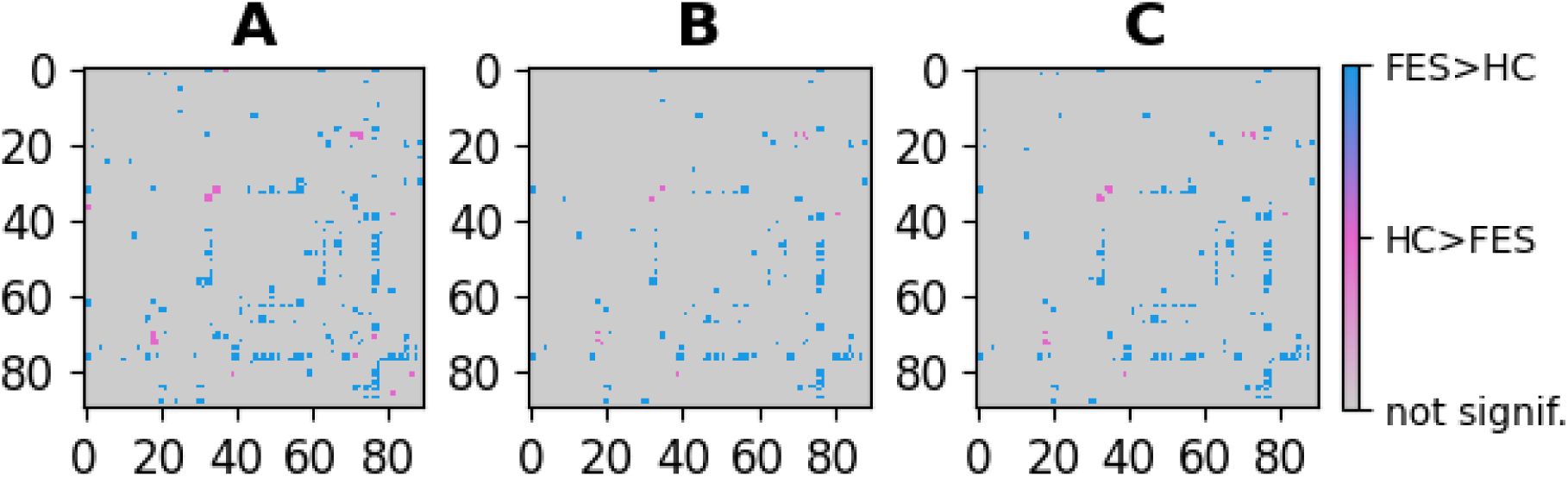
Difference between HC and FES. The significant differences after the p<0.05 FDR correction are shown in blue (higher FC in patients) and pink (higher FC in healthy controls). A) Default analysis; B) Analysis including additional inter-subject correction for the amount of motion; C) Analysis including additional inter-subject correction for the amount of motion, as well as rejecting outliers. See the Discussion section for details of the additional analysis shown in the B and C panels.

### 3.2 Association between symptom severity, medication, duration of untreated psychosis, duration of antipsychotic treatment, and functional connectivity

We have observed a significant correlation between symptom severity measured by PANSS and functional connectivity, see Figure 3. To increase the power by focusing on the most relevant candidate connections, we used a masking matrix of p-values from a group comparison of healthy volunteers and patients to distinguish regions with significantly greater functional connectivity in patients. PANSS positive scale significantly correlated with the functional connectivity between the right precentral gyrus and the right middle cingulate gyrus. PANSS negative scale significantly correlated with the functional connectivity between the right middle cingulate gyrus and the left lingual gyrus.

**Figure 3:**
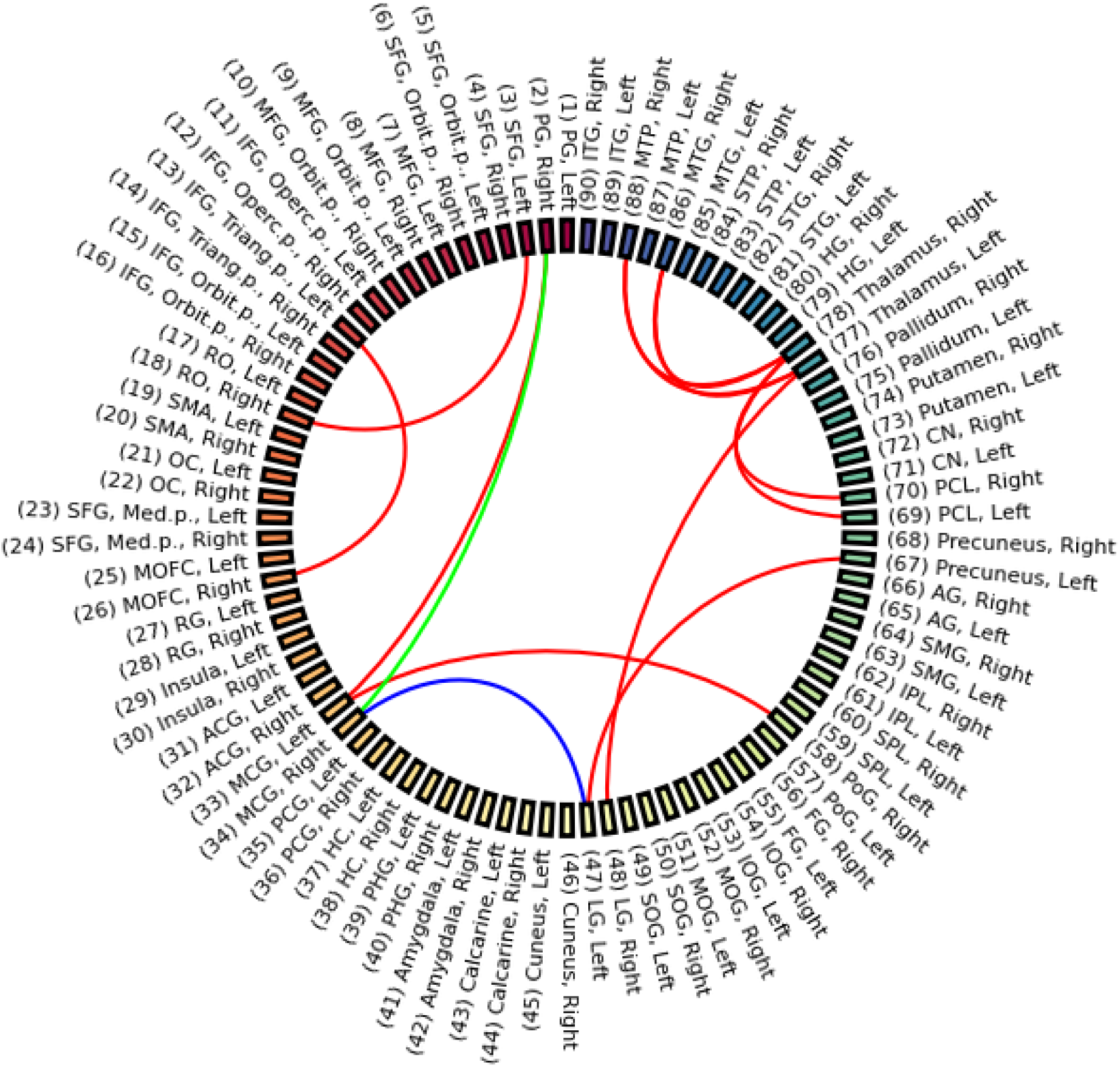
Association between antipsychotic medication, symptom severity, and functional connectivity. Antipsychotic medication significantly correlated with the functional connectivity of 12 ROI pairs (red links), PANSS Positive scale significantly correlated with the functional connectivity of one ROI pair (green link), and PANSS Negative scale significantly correlated with the functional connectivity of one ROI pair (blue link). Results with p*<*0.05 (FDR-corrected) are shown.

Further, we have observed a significant correlation between antipsychotic medication and functional connectivity. Regions with the most links included the bilateral thalamus, left middle cingulate gyrus, right middle temporal gyrus, and right middle temporal pole; see Table 1 for the full list of connections.

**Table 1:** Correlation between antipsychotic medication, PANSS Positive, PANSS Negative, and functional connectivity in patients; note that only p*<*0.05 (FDR-corrected) are reported.

| Region 1 | Region 2 | Spearman r |
| --- | --- | --- |
| ***Antipsychotic medication*** |  |  |
| Thalamus, Left | Middle Temporal Pole, Right | 0.33 |
| Superior Frontal Gyrus, Left | Rolandic Operculum, Right | 0.32 |
| Thalamus, Right | Middle Temporal Gyrus, Right | 0.31 |
| Thalamus, Right | Middle Temporal Pole, Right | 0.31 |
| Inferior Frontal Gyrus, Triangular Part, Left | Medial Orbital Frontal Cortex, Right | 0.31 |
| Lingual Gyrus, Right | Thalamus, Left | 0.30 |
| Lingual Gyrus, Left | Precuneus, Left | 0.30 |
| Paracentral Lobule, Left | Thalamus, Right | 0.29 |
| Precentral Gyrus, Right | Middle Cingulate Gyrus, Left | 0.29 |
| Thalamus, Left | Middle Temporal Gyrus, Right | 0.27 |
| Middle Cingulate Gyrus, Left | Postcentral Gyrus, Right | 0.27 |
| Paracentral Lobule, Right | Thalamus, Right | 0.27 |
| ***PANSS Positive*** |  |  |
| Precentral Gyrus, Right | Middle Cingulate Gyrus, Right | 0.35 |
| ***PANSS Negative*** |  |  |
| Middle Cingulate Gyrus, Right | Lingual Gyrus, Left | 0.33 |

We have also observed a significant correlation between antipsychotic medication and symptom severity measured by PANSS Total (Spearman r=0.33, p*<*0.001).

Conversely, we found no relationship between the duration of untreated psychosis, duration of antipsychotic treatment, and functional connectivity.

### 3.3 Effect of data-denoising on functional connectivity

To evaluate the stability of the above-presented findings with respect to the strictness of denoising, we also performed our analysis with moderate and raw denoising settings. In the moderate denoising, healthy volunteers exhibited significantly higher (p*<*0.05 FDR-corrected) functional connectivity in 1098 ROI-pairs compared to patients. Conversely, 43 ROI pairs had significantly higher (p*<*0.05 FDR-corrected) functional connectivity in patients, see Figure 5B. The most increased links in patients included the bilateral thalamus, bilateral superior temporal gyrus, and left postcentral gyrus (see Table S3 for the full list of affected connections). In the opposite direction, the most decreased links in patients involved the right precentral gyrus, bilateral postcentral gyrus, left rolandic operculum, and the right paracentral lobule (see Table S4 for the full list of affected connections).

In stark contrast to the stringent denoising scheme, the median functional connectivity was greater in patients for the moderate denoising scheme. The median connectivity difference between healthy controls and patients in the moderate scheme was 0.0349 (median FC in patients=0.1388, MAD=0.0338, median FC in controls=0.1737, MAD=0.0477, u=5762 (z-score=-3.1351), p<0.001). Indeed, the disease effect on functional connectivity indices was clearly biased towards hypoconnectivity in the moderately denoised dataset, while there was a relatively balanced hypoconnectivity and hyperconnectivity (with a slight prevalence of hyperconnectivity) observed when using the default stringent denoising scheme, see Figure 4C.

**Figure 4:**
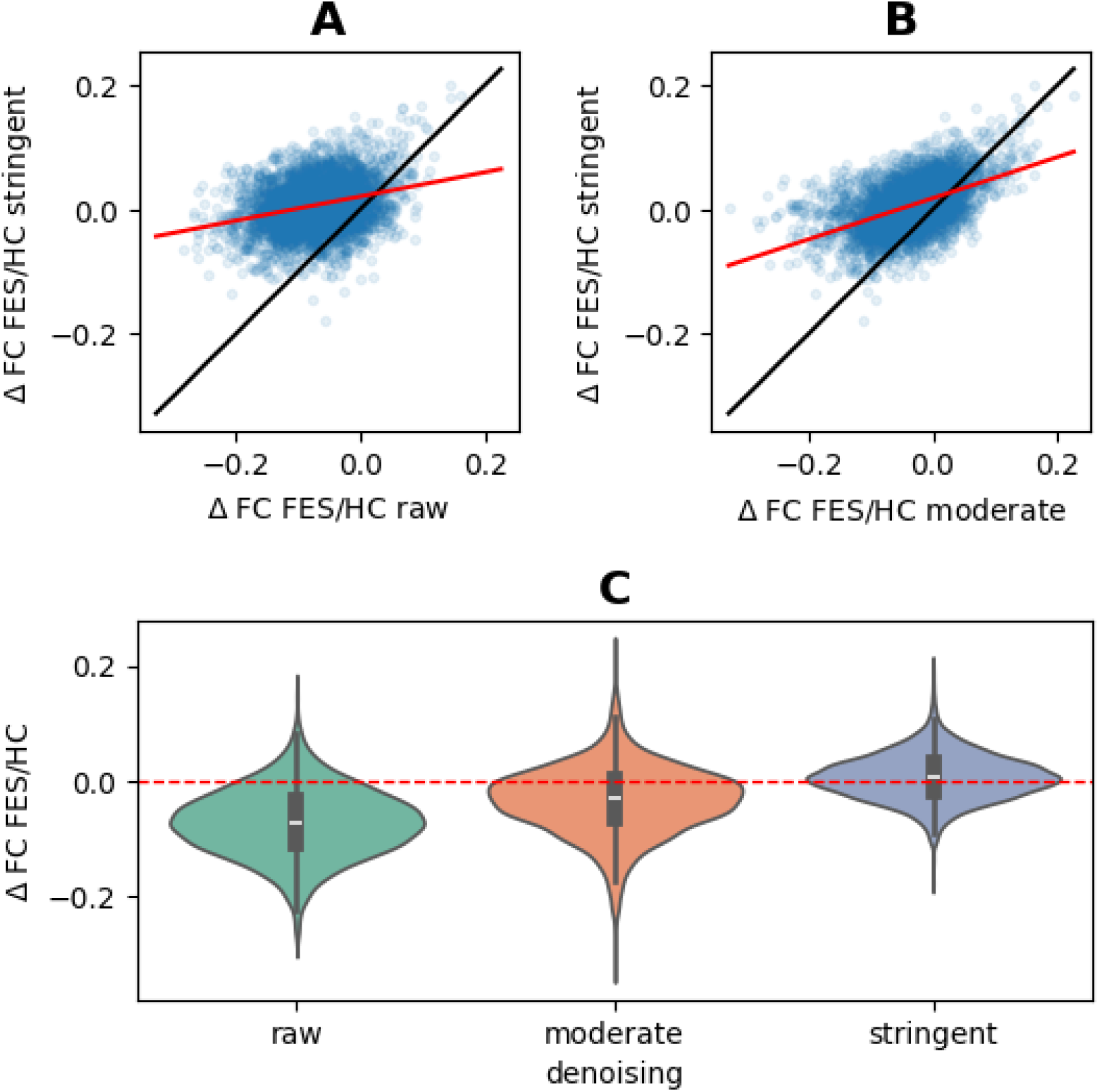
Comparison of the effect of disease on functional connectivity observed with the raw, moderate, and stringent denoising. A) Scatterplot of the median FC difference between patients and healthy controls in the raw and stringent scheme. B) Scatterplot of the median FC difference between patients and healthy controls in the moderate and stringent scheme. Black lines mark identity, and the red line corresponds to the best linear fit. Each dot corresponds to one pair of regions, showing the difference in their medians in the two denoising schemes. C) Violin plot of the FC differences between healthy controls and patients in the raw, moderate, and stringent denoising schemes.

In the case of moderate denoising, we found a significant correlation between functional connectivity and antipsychotic medication. Regions included the bilateral thalamus, bilateral rolandic operculum, and bilateral lingual gyrus (see Table S5 for the full list of affected connections).

In the raw denoising, healthy volunteers exhibited significantly higher (p*<*0.05 FDR-corrected) functional connectivity in 1281 ROI-pairs compared to patients; see Table S6 for the full list of affected connections. Conversely, no ROI pairs had significantly higher (p*<*0.05 FDR-corrected) functional connectivity in patients, see Figure 5A. The median connectivity difference between healthy controls and patients in the raw scheme was 0.0611 (median FC in patients=0.5025, MAD=0.1132, median FC in controls=0.5636, MAD=0.1190, u=5509 (z-score=-2.4224), p<0.01). The disease effect on functional connectivity indices was thus clearly biased towards hypoconnectivity in the raw scheme, see Figure 4C. We also observed no significant relation between functional connectivity and any of the clinical variables.

**Figure 5:**
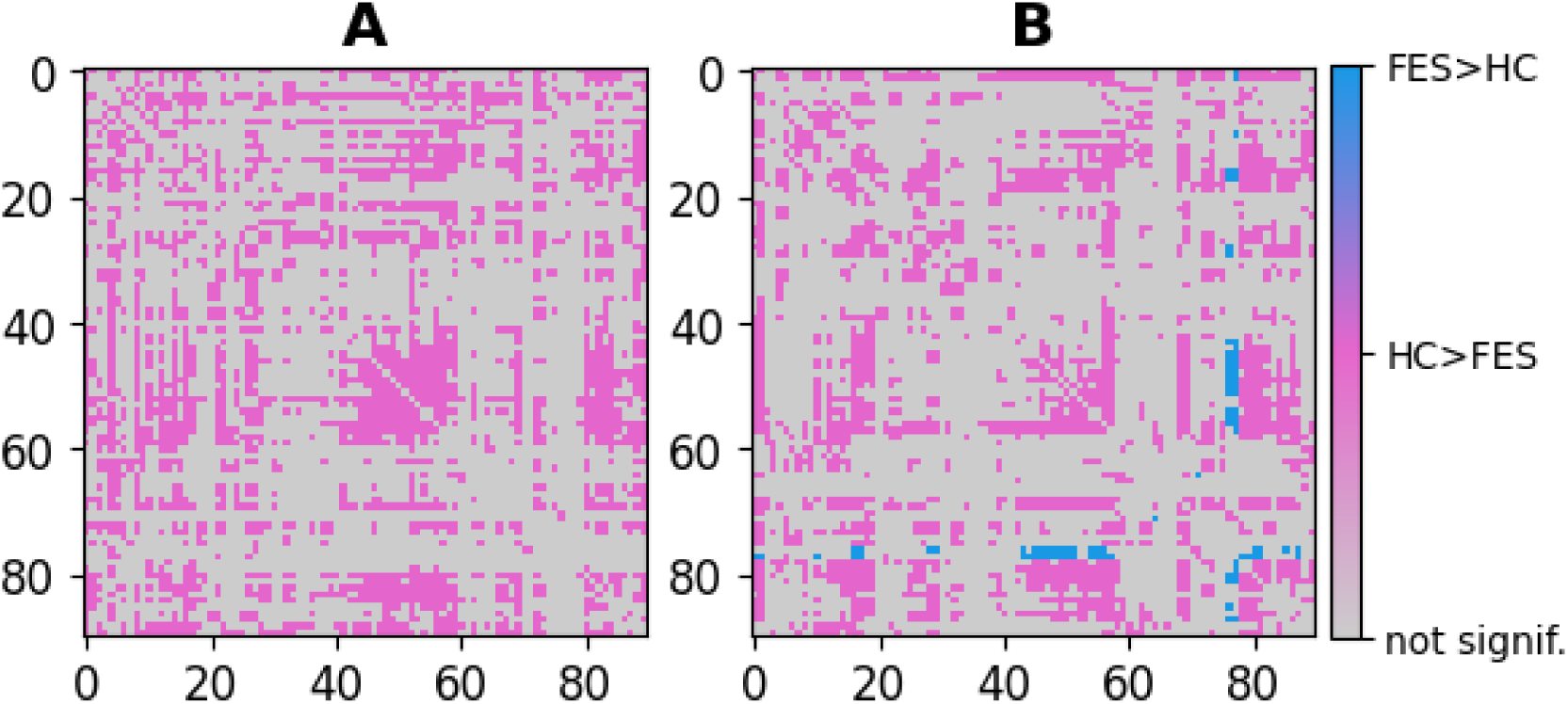
Difference between HC and FES for the A) raw and B) moderate denoising variants. The differences significant after the p*<*0.05 FDR correction are shown in blue (higher FC in patients) and pink (higher FC in healthy controls).

Despite these vast differences in the overall shift of FC between the two populations, the spatial localization of the disease effect observed with the three denoising schemes was strongly correlated (raw and stringent: Spearman r=0.25, p<0.001; moderate and stringent: Spearman r=0.48, p<0.001), see Figure 4A and 4B respectively for the scatterplot across region pairs.

## 4 Discussion

Using the stringent denoising scheme, we observed that individuals with FES showed a pattern of focal hyperconnectivity in thalamic, basal ganglia, and cingulo-sensorimotor circuits, combined with cortical hypoconnectivity. This profile aligns with models of schizophrenia that conceptualize early psychosis as a disturbance of hierarchical cortico–subcortical integration, with thalamic and striatal systems exhibiting increased (possibly compensatory) reactivity against a background of impaired cortical coordination (Anticevic et al. 2015; Wengler et al. 2020). This hybrid pattern replicates prior work documenting thalamo–sensorimotor hyperconnectivity in schizophrenia (Anticevic et al. 2015; Woodward et al. 2012) and suggests an early imbalance in hierarchical predictive processing.

The most increased links in patients involved the bilateral thalamus, globus pallidus, bilateral middle cingulate gyrus, bilateral supramarginal gyrus, right superior occipital gyrus, and bilateral precuneus. These findings mirror the “thalamocortical dysconnectivity” framework proposed by Anticevic et al. (2014) and the results of meta-analyses reporting increased thalamic coupling to sensorimotor cortices in psychosis. According to these models, thalamic hyperconnectivity may reflect insufficient top-down regulation from prefrontal and hippocampal systems, leading subcortical hubs to over-signal or over-synchronize with cortical targets. Notably, a subset of the hyperconnected edges identified here was also implicated in symptom severity and medication effects, suggesting that these subcortical–cingular–motor circuits may represent a core network phenotype of early schizophrenia.

In contrast, the most decreased links in patients involved circuits linking the supplementary motor area and precentral cortex with the hippocampus, caudate, and putamen. This pattern is consistent with meta-analytic evidence for widespread hypoconnectivity across fronto-parietal, hippocampal, and cortico-striatal networks in schizophrenia (Dong et al. 2018; S. Li et al. 2019). These networks support predictive processing, contextual integration of prior information, and precision-weighted gating of sensory signals (Friston 2005; Corlett et al. 2019). Their reduced functional integration in FES aligns with computational theories proposing that weakened cortical priors and impaired hippocampal-prefrontal coordination degrade the system’s ability to maintain stable internal models and suppress noisy bottom-up prediction errors (Sterzer et al. 2018; Sigurdsson & Duvarci 2016). In this context, thalamic and striatal hyperconnectivity may arise as maladaptive attempts to stabilize cortical dynamics in the face of insufficient cortico-hippocampal control, a view that is compatible with theoretical work showing that hierarchical heterogeneity of cortical microcircuits shapes large-scale neural dynamics and stability (Demirtaş et al. 2019). Our findings also align with longitudinal evidence in antipsychotic-naive FES showing widespread baseline dysconnectivity and that antipsychotic exposure in antipsychotic-naive FES is associated with increased functional connectivity primarily between the thalamus and the rest of the brain (Chopra et al., 2021). A striking and unifying observation across all analyses was the role of the mid-cingulate cortex (MCC). MCC appeared as a major hyperconnected hub in patients, but also as the key node associated with both PANSS symptom dimensions and CPZ-equivalent antipsychotic exposure (see below). This convergence suggests MCC as the principal clinically relevant region in our dataset. The MCC is an integrator of salience, motor preparation, effort allocation, and executive control, functions that are highly disrupted in psychosis. Its involvement here strongly supports theories emphasizing abnormalities in salience signaling and motor relevance attribution in schizophrenia (Shackman et al. 2011; Hoffstaedter et al. 2014; Kapur 2003). This is further supported by studies demonstrating aberrant recruitment of the medial cingulate/ACC region during cognitive control tasks in schizophrenia during cognitive-control and motor-preparatory tasks in psychosis (Heckers et al. 2004).

Furthermore, brain hyperconnectivity correlated with both symptom severity as well as antipsychotic medication in subjects at their first episode of psychosis. The positive symptom dimension (PANSS Positive) was associated with FC between right precentral gyrus and right MCC, indicating enhanced coupling between motor cortex and a core salience/motor-control hub (Shackman et al. 2011; Du et al. 2019; Kapur 2003). This pattern is highly compatible with Kapur’s aberrant salience model and with contemporary models linking psychosis to dysregulated interactions between salience networks and motor/output systems. Conversely, the negative symptom dimension (PANSS Negative) was associated with functional connectivity between the right MCC and the left lingual gyrus in our sample, suggesting altered coupling between a cingulate salience/control hub and visual–perceptual cortex. While this specific MCC–lingual link is not a canonical finding, it is consistent with broader evidence that negative symptom dimensions relate to salience-network dysfunction and abnormal salience–default mode (and salience–sensory) communication in schizophrenia (Hare et al. 2019; Palaniyappan & Liddle 2012). These findings are consistent with the literature suggesting that negative symptom dimensions are associated with cingulate/salience-network dysfunction and with altered coupling between salience/control systems and sensory–perceptual networks, potentially contributing to impaired contextual appraisal.

This pattern most likely reflects a combination of illness severity, given that CPZ-equivalent dose is clinically titrated and therefore is often confounded with positive symptom severity in naturalistic samples (Troisi et al. 1997; Ringen et al. 2019), and medication-driven modulation of salience-related motor circuitry, in line with evidence showing antipsychotic effects on thalamo-cortical and cingulate networks (Chopra et al. 2021; Sarpal et al. 2015). Importantly, Chopra et al. demonstrated that DMN–limbic normalization can occur even without medication, whereas drug exposure specifically alters thalamo-cingulate circuits. Our findings extend this by showing that in a naturalistic FES sample, antipsychotic exposure was most strongly associated with the same motor–salience connections that also tracked positive symptom severity in our data. The overlap between CPZ-related and PANSS Positive-related connectivity further supports the interpretation that apparent medication effects may partly reflect underlying symptom burden.

It is worth mentioning some technical considerations of the present study. Several denoising strategies were discussed in recent papers. While a unique consensus has not been reached in the field, the implementation of a more stringent denoising scheme has become a standard practice (Power et al. 2014; Patel et al. 2014), motivating our default choice. To quantify functional connectivity, in line with common practice, we use Pearson’s linear correlation coefficient, as it was shown to be sufficient under standard conditions, as with the current dataset (Hlinka et al. 2011). A possibly fruitful further direction would indeed lie in extension to mapping the consequences or analogies of our findings to other network characteristics previously shown to be altered in clinical high-risk and early-illness schizophrenia individuals, in particular to dynamic functional connectivity (Mennigen et al. 2019), and amplitude of low frequency fluctuations or graph-theoretical functional dysconnectivity characterization (Tang et al. 2019). On the other side, utilizing the full FC matrix has shown an advantage in machine learning classification of schizophrenia above and beyond some advanced methods such as topological data analysis (Caputi et al. 2021).

There was a considerable difference between the stringent, moderate, and raw denoising schemes in the number of significant ROI pairs in each group comparison. Both raw and moderate denoising schemes resulted in a higher number of ROI pairs that showed significantly higher FC in healthy controls compared to patients (1281/0 in raw and 1098/43 in moderate), while in the stringent denoising scheme, the proportion changed as we observed a slightly higher number of ROI pairs exhibiting hyperconnectivity in patients (15/150). Note that the use of a substantially stricter denoising scheme than used in a range of studies in the literature may explain some of the deviation of our results with respect to the field. Despite the differences between denoising strategies, median connectivity matrices are strongly correlated within each group (the correlation of FC between raw and stringent was 0.5301 (p<0.001) in healthy individuals and 0.5681 (p<0.001) in patients, and between moderate and stringent was 0.8798 (p<0.001) in healthy individuals and 0.9181 (p<0.001) in patients. Also, the difference between the groups, i.e., the disease effect, is strongly correlated (across region pairs) between the denoising strategies (raw and stringent: Spearman r=0.25, p<0.001; moderate and stringent: Spearman r=0.48, p<0.001), see Figure 4A and 4B, respectively, for the scatterplot across region pairs. Although we found overlapping ROI-pairs between the stringent and moderate denoising that exhibited correlation with antipsychotic medication (namely, regions involving the bilateral thalamus, right middle temporal pole, right middle temporal gyrus, and right lingual gyrus), we found no relationship between functional connectivity and any of the selected clinical variables when the raw data were used, confirming that minimal standard preprocessing is needed to reduce noise in the data.

To ascertain that the results were not driven by any residual artifacts due to differences in head motion, we also carried out a control analysis where mean framewise displacement (FD), the aggregated measure of subject motion, was included as a covariate of no interest. The results again remained qualitatively equivalent, suggesting that the effect of motion on FC is relatively small compared to the effect of disease; see Figure 2B. The results remained largely unchanged after the additional removal of the 14 subjects who showed maximum head motion larger than 1.5×IQR (interquartile range), the Tukey criterion for detection of outliers (Tukey 1977); see Figure 2C. The difference in the number of ROI pairs with significantly changed FC could be due to lower degrees of freedom in the additional corrected analyses.

## 5 Conclusions

Our study using a whole-brain functional connectivity approach on a relatively large sample has shown a relatively balanced, yet spatially resolved, picture of hyperconnectivity and hypoconnectivity in first-episode psychosis while demonstrating a significant association between the affected connection strengths and clinical symptoms and cumulative dosage of antipsychotic medication. Our results support a model in which early schizophrenia combines cortical hypoconnectivity with subcortical and cingulo-sensorimotor hyperconnectivity, forming a hybrid dysconnectivity profile that is closely tied to symptoms and treatment. Importantly, we have shown that this balanced picture gets substantially skewed in the direction of dominantly observed hypo- or dysconnectivity when applying more moderate data-denoising schemes, while the relation to symptoms is rendered insignificant. While the results of all approaches are in overall qualitative agreement, including with localization of most commonly reported disease connectivity effects in the literature, taking into account the results obtained with the current stringent denoising may provide a stronger relation to clinical variables and contribute to the understanding of the diversity of previously reported results, as well as the interpretation of schizophrenia as a dysconnection disease in general.

## 6 Acknowledgments and Disclosures

The publication was supported by: ERDF-Project Brain dynamics, No. CZ.02.01.01/00/22_008/000464, Czech Health Research Council Project No. NU21-08-00432 and by the MSMT CR program Cooperatio, Neuroscience, Charles University.

This manuscript has been posted on a preprint server, BioRxiv, under doi: https://doi.org/10.1101/2024.09.20.613853.

## Author contributions

FS, JH, and JHl are responsible for conceptualization; DT and JHl are responsible for methodology; DT is responsible for software; DT a JHl performed the validation; DT, JH, and JHl are responsible for the formal analysis; DT, MK, BRB, FS, JH, and JHl are responsible for investigation; JT, FS, and JHl provided the resources; DT, JT, FS, JH, and JHl are responsible for data curation; DT, MK, BRB, JH, and JHl wrote the original draft; DT, MK, BRB, JT, FS, JH, and JHl reviewed and edited the manuscript; DT is responsible for the visualization; JT, FS, JH, and JHl performed the supervision; JT, FS, and JHl administered the project; FS, JH and JHl are responsible for funding acquisition.

## Data Availability Statement

The data that support the findings of this study are available from the corresponding author upon reasonable request.

